# Splicing of the SynGAP Carboxyl-Terminus Enables Isoform-Specific Tuning of NMDA Receptor Signaling Linked to Cognitive Function

**DOI:** 10.1101/2020.01.18.911487

**Authors:** Murat Kilinc, Thomas K. Creson, Camilo Rojas, Sabyasachi Maity, Aliza A. Le, Julie Lauterborn, Brent Wilkinson, Nicolas Hartel, Nicholas Graham, Adrian Reich, Gemma Gou, Yoichi Araki, Àlex Bayés, Marcelo P. Coba, Gary Lynch, Courtney A. Miller, Gavin Rumbaugh

**Affiliations:** Graduate School of Chemical and Biological Sciences, The Scripps Research Institute, The Scripps Research Institute, Jupiter, FL, USA; Departments of Neuroscience and Molecular Medicine, The Scripps Research Institute, The Scripps Research Institute, Jupiter, FL, USA; Bioinformatics and Statistics Core, The Scripps Research Institute, The Scripps Research Institute, Jupiter, FL, USA; Department of Anatomy and Neurobiology, The University of California, Irvine, CA, USA; Zilkha Neurogenetic Institute, Keck School of Medicine, University of Southern California, Los Angeles, CA, USA; Mork Family Department of Chemical Engineering and Materials Science, University of Southern California, Los Angeles, California; Molecular Physiology of the Synapse Laboratory, Biomedical Research Institute Sant Pau (IIB Sant Pau), Barcelona, Spain; Universitat Autònoma de Barcelona, 08193 Bellaterra (Cerdanyola del Vallès), Spain; Department of Neuroscience, Johns Hopkins University School of Medicine, Baltimore, MD 21205, USA

**Keywords:** *Syngap1*, SynGAP, Synapse, PSD95, PDZ domain, Long-term potentiation, Intellectual disability, Autism spectrum disorder, Epilepsy

## Abstract

SynGAP-α1 is a splice variant of the neurodevelopmental disorder risk gene, *SYNGAP1/Syngap1*. α1 encodes the C-terminal PDZ binding motif (PBM) that promotes liquid-liquid phase separation, a candidate process for postsynaptic density organization within excitatory synapses. However, it remains unknown how the endogenous SynGAP PBM regulates synapse properties and related cognitive functions. We found that a major PBM function in mice is to limit the mobility of SynGAP-α1 in response to NMDA receptor activation. Genetic disruption of the PBM increased SynGAP-α1 mobility to levels consistent with other non-PBM-containing C-terminal isoforms. This resulted in a lowering of the threshold for NMDA receptor-dependent signaling required for plasticity, leading to aberrant strengthening of excitatory synapses in spontaneously active neurons. PBM-deficient animals also exhibited a lower seizure threshold, disrupted LTP, and impaired cognition. Thus, the PBM enables isoform-specific SynGAP gating of NMDA receptor function, a mechanism linking synaptic signaling dynamics to network excitability and cognition.

## Introduction

Higher cognitive processes, such as learning and memory, are thought to rely on dynamic regulation of excitatory synapse structure and function (Hayashi-Takagi et al., 2015; Penn et al., 2017). The postsynaptic density (PSD) is an electron-dense protein complex that resides within dendritic spines and is a structure required for dynamic regulation of excitatory synapse function (Kennedy et al., 2005). A function of the PSD is to transduce synaptic NMDAR opening and calcium influx into the eventual capture of AMPARs at the post-synaptic terminal, which is an effector mechanism that enables the expression of a major form of synaptic plasticity linked to learning (Huganir and Nicoll, 2013; Kessels and Malinow, 2009). This has led to the idea that the organization and proper functioning of the PSD is required for higher cognitive functions and alterations to PSD-related complexes contributes to disease (Grant, 2012). This idea is supported by genetic evidence. Indeed, genes that encode PSD proteins are enriched for pathological variants in patient cohorts defined by impaired cognitive function (Bayes et al., 2014; Bayes et al., 2011; Kirov et al., 2012). However, it remains unclear how the PSD forms and subsequently carries out these dynamic functions. The PSD contains unusually high levels of PDZ domain-containing proteins, as well as related proteins that contain PDZ binding motifs (PBMs) (Feng and Zhang, 2009). These types of proteins are thought to organize the PSD through a form of liquid-liquid phase separation (LLPS) (Zeng et al., 2016).

Pathogenic variants in *SYNGAP1/Syngap1*, the gene encoding SynGAP protein, are a leading cause of sporadic neurodevelopmental disorders (NDDs) defined by impaired cognitive function, epilepsy, and autistic features (Deciphering Developmental Disorders, 2015, 2017; Hamdan et al., 2009; Mignot et al., 2016; O’Roak et al., 2014; Parker et al., 2015; Satterstrom et al., 2019; Vlaskamp et al., 2019). SynGAP contains a PBM that promotes LLPS when binding to PSD95-family proteins (Zeng et al., 2016). Studying the SynGAP PBM is of considerable interest because it is one of the most abundant proteins in the PSD, where it is expressed at roughly equivalent levels compared to PSD95. Moreover, studying the molecular features of SynGAP may provide insight into the mechanisms of severe NDDs. *In vitro* studies have suggested that the PBM is important for regulating synapse structure and function (Rumbaugh et al., 2006; Vazquez et al., 2004). However, the contribution of the endogenous SynGAP PBM to PSD assembly and/or signaling dynamics remains unknown because its function has not been studied in isolation nor in the context of normal SynGAP protein expression. This is critical because *Syngap1* encodes several alternatively spliced protein isoforms, with only a subset expressing a PBM (Chen et al., 1998; Kim et al., 1998; McMahon et al., 2012).

The SynGAP PBM arises from alternative splicing within the last exon of *Syngap1* into the Alpha 1 (α1) reading frame. This leads to the expression of a unique 12 amino acid, PBM-containing C-terminus (Chen et al., 1998; Kim et al., 1998). Thus, to precisely understand the functions of the SynGAP PBM, this motif must be selectively targeted at the genomic level so that the expression and coding sequences of the other SynGAP isoforms remain unperturbed. By doing this, it is theoretically possible to dissociate the precise role of the PBM from other known SynGAP functions. Indeed, other C-terminal, non-PBM-containing SynGAP isoforms, such as β and α2, have alternative C-terminal sequences that result in unique protein-protein interactions, synaptic targeting profiles, and/or functional outcomes within dendritic spines (Li et al., 2001; McMahon et al., 2012; Moon et al., 2008). Here, we created a mouse model expressing the endogenous SynGAP-α1 isoforms with point mutations in the PBM that disrupted binding to PDZ domains. The model revealed that a major function of this PBM is to limit the mobility of SynGAP-α1 in response to NMDAR activity. Disrupting the PBM lowered the threshold for NMDAR-mediated signaling required for synaptic plasticity. This synaptic deficit was linked to aberrantly strong excitatory synapses, altered synaptic plasticity, reduced seizure threshold, and impaired cognitive function. We conclude that the PBM within SynGAP-α1 is crucial for determining the levels of NMDAR activity required for synaptic potentiation. This postsynaptic mechanism, enabled through *Syngap1* C-terminal splicing, may promote memory-related cognitive functions and balanced excitability within distributed hippocampal/cortical networks.

## Results

Our goal was to create a mouse line with a disrupted PBM without altering other SynGAP isoforms. Before doing this, we had to identify PBM-disrupting point mutations within the α1 coding sequence that were silent within the open reading frames of the remaining C-terminal isoforms. *In silico* predictions and prior studies (Rumbaugh et al., 2006; Zeng et al., 2016) suggested that a double point mutation within the α1 PBM could meet these requirements (**Fig. 1A-B)**. To test this, we introduced the point mutations into cDNAs and tested their impact on PDZ binding. Using an established cell-based assay that reports PDZ binding between the SynGAP PBM and PSD95 (Zeng et al., 2016), we found that these point mutations dramatically disrupted SynGAP-PDZ binding. When expressed individually in HeLa cells, PSD95-tRFP localized to the cytoplasm, while a SynGAP fragment containing the coiled-coil domain and α1 C-tail (EGFP-CCα1) was enriched in the nucleus **(Fig. 1C-E)**. The co-expression of these two proteins led to SynGAP localization into the cytoplasm. However, this shift in localization did not occur when PBM point mutations were present **(Fig. 1D-E)**, indicating that the selected amino acid substitutions impaired binding to the PDZ domains. Moreover, co-immunoprecipitation in heterologous cells indicated that the point mutations in the PBM disrupted the direct association of full-length SynGAP-α1 with PSD95 (Fig. S1A-B). Finally, these point mutations resulted in reduced synaptic enrichment of exogenously expressed SynGAP fragments in cultured forebrain neurons (Fig. S1C-E). Based on this evidence, we introduced the point mutations into the mouse *Syngap1* gene through homologous recombination **(Fig. 1A,F-H)**. Both heterozygous and homozygous PBM mutant animals (hereafter *Syngap1*^*+/PBM*^ or *Syngap1*^*PBM/PBM*^) were viable with no obvious gross morphological features. We observed Mendelian ratios after interbreeding *Syngap1*^*+/PBM*^ animals (Fig. S1F), indicating that disrupting the PBM did not impact survival. Western blot analysis of forebrain homogenates isolated from *Syngap1*^*+/PBM*^ or *Syngap1*^*PBM/PBM*^ mutant animals demonstrated no difference in total SynGAP (t-SynGAP) protein levels using antibodies that detect all SynGAP splice variants **(Fig. 1I-J)**. Moreover, using isoform-selective antibodies (Gou et al., 2019), we observed normal expression of SynGAP-β and SynGAP-α2 isoforms **(Fig. 1I-J)**. A reduced signal of ∼60% was observed in samples probed with α1-specific antibodies. However, we observed a similarly reduced signal in heterologous cells expressing a cDNA encoding the mutant PBM (Fig. S1G-I), demonstrating these antibodies have reduced affinity for mutated α1 spliced sequence containing the altered PBM. These data indicate that the α1 variant is expressed normally in *Syngap1*^*PBM/PBM*^ animals. This was supported by RNA-seq data, where normal levels of mRNA containing the α1 reading frame were observed (Fig. S1J). These data, combined with the observation of no change in total SynGAP protein expression in *Syngap1*^*PBM/PBM*^ samples **(Fig. 1I-J)**, strongly support the conclusion that the PBM-disrupting point mutations do not change the expression levels of the major SynGAP C-terminal splice variants, including those containing the PBM. Thus, this animal model is suitable for studying the biological functions of endogenous SynGAP PBM binding.

**Figure 1.**
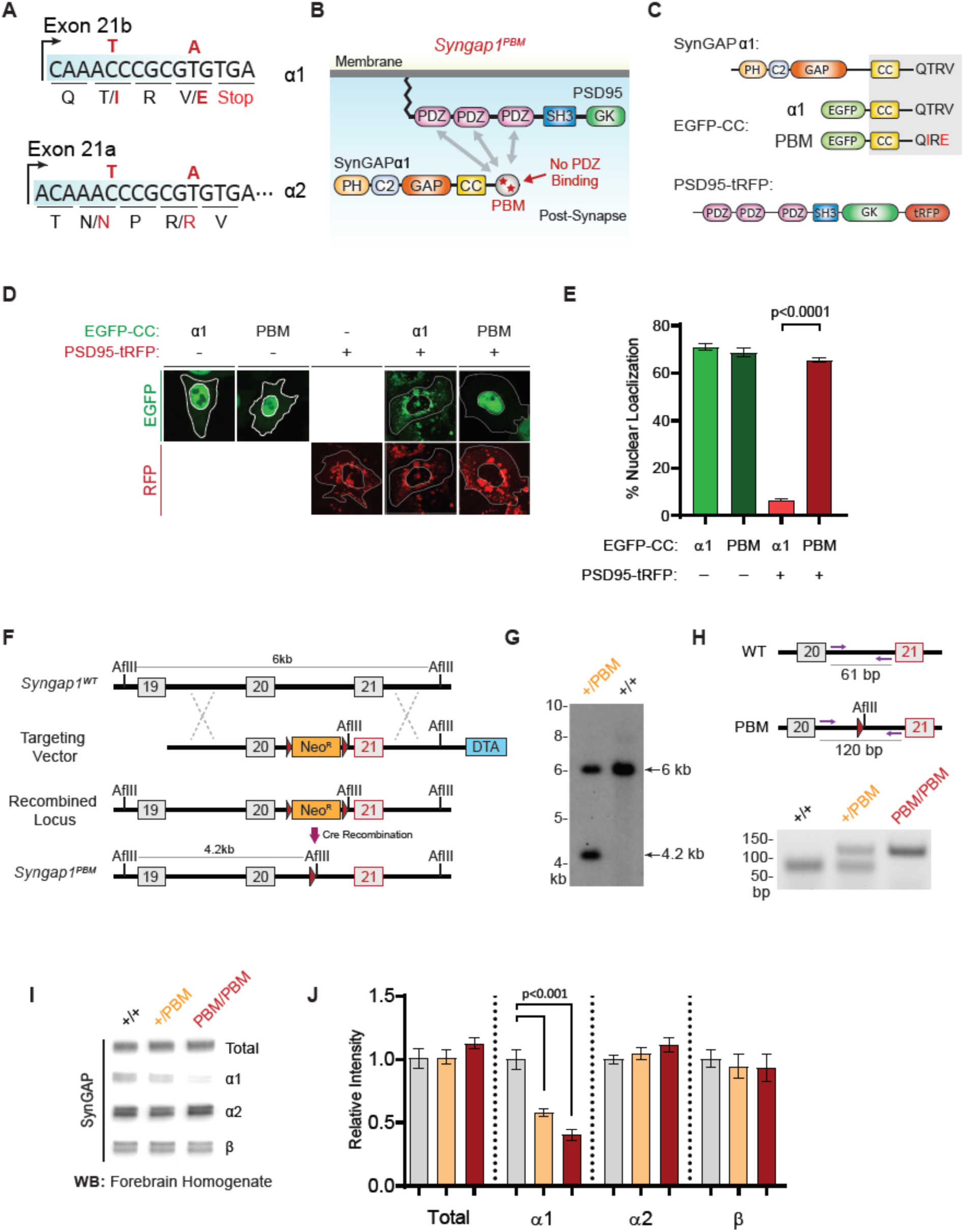
Targeted genomic disruption of SynGAP-α1 PDZ-binding through introduction of missense point mutations. **(A)** Schematic diagram for exon map and alternative use of exon 21 in the *Syngap1* gene. Exon 21b encodes for α1 isoform. exon 21a encodes for α2 isoform. Point mutations indicated in red alter exon 21b coding sequence without influencing exon21a open reading frame. **(B)** Schematics of SynGAP-α1 and PSD95 domain structure and the location of point mutations. **(C)** Illustrations of constructs expressed in HeLa cells to study PDZ-dependent interaction between SynGAP and PSD95. EGFP-CC constructs are homologous to SynGAP-α1 C-terminus. **(D)** Co-localization of EGFP-CCα1 and PSD95-tRFP in HeLa Cells. Representative images showing subcellular localizations of WT (α1) or PDZ-binding mutant (PBM) EGFP-CCα1 and PSD95-tRFP in HeLa cells when expressed individually or together. **(E)** Quantification of D. ANOVA with Tukey’s multiple comparisons test, F(3, 96) = 531.4, p<0.0001, n=23-25 cells per condition. **(F)** Schematics of the targeting strategy. The targeting vector was spanning Exon20 & 21. The vector included point mutations in Exon21, a neomycin resistance selection cassette flanked by Cre recombination sites and diphtheria toxin selection cassette (DTA). **(G)** Southern blot analysis showing the genomic DNA of the tested heterozygous mice compared to C57BL/6J wild-type DNA following AflII digestion and hybridization with external 5’ probe spanning exon 19. **(H)** PCR based genotyping strategy. Primers flanking leftover LoxP site yields 61bp product in WT and 120bp product in mutated allele. **(I)** Representative western blots showing expression levels of total SynGAP and individual isoforms in forebrain lysates. **(J)** Quantification of I. Relative intensity of bands normalized to total protein signal. ANOVA with Tukey’s multiple comparisons test. SynGAP-α1: F(2, 14) = 24.86, p<0.0001; +/+ vs +/PBM: p=0.0009; +/+ vs PBM/PBM: p<0.0001. n=5 per genotype.

SynGAP is a major regulator of neuronal excitability and reduced SynGAP function leads to larger and stronger excitatory synapses (Kilinc et al., 2018). Therefore, we reasoned that PBM mutations would impact excitatory synapse structure and function in neurons from *Syngap1*^*PBM/PBM*^ animals. In forebrain neurons cultured from these mice, we observed larger dendritic spines with increased surface AMPA receptor (GluA1) expression **(Fig. 2A)**. In acute hippocampal slices, there was an increase in synaptic excitability **(Fig. 2B)** and *in vivo*, we detected a reduced seizure threshold in *Syngap1*^*PBM/PBM*^ mice **(Fig. 2C)**. Each of these phenotypes are consistent with past studies that probed neurons with reduced SynGAP function (Clement et al., 2012; Rumbaugh et al., 2006; Vazquez et al., 2004). However, given that steady state SynGAP expression is normal in these mice **(Fig. 1I-J)**, we hypothesized that PBM mutations may disrupt a biochemical function of SynGAP within excitatory synapses. SynGAP is known to regulate dendritic spine Ras/ERK signaling through dynamic changes in its expression within the PSD (Araki et al., 2015; Rumbaugh et al., 2006). Thus, we were interested in determining how disrupting the PBM would impact SynGAP levels in the synapse. t-SynGAP levels were reduced in PSD fractions prepared from the hippocampus of *Syngap1*^*PBM/PBM*^ mice **(Fig. 2D,E)**. Importantly, a corresponding increase in t-SynGAP was observed in the triton soluble synaptosomal fraction in these mice **(Fig. 2F)**, further supporting the observation of reduced t-SynGAP levels in the PSD. We observed similar reductions in t-SynGAP levels within the PSD **(Fig. 2G,H)** and dendritic spines (Fig. S2A,B) of primary cultured neurons from *Syngap1*^*PBM/PBM*^ mice. Consistent with reduced SynGAP PSD levels (Araki et al., 2015), ERK1/2 signaling was elevated in neurons cultured from *Syngap1*^*PBM/PBM*^ mice **(Fig. 2I)**. We next explored the mechanisms underlying changes in synaptic properties in *Syngap1*^*PBM/PBM*^ mice. Acute treatment with the NMDAR antagonist APV normalized SynGAP levels in both PSD preparations **(Fig. 2J,K)** and dendritic spines (Fig. S2A,C) in neurons derived from *Syngap1*^*PBM/PBM*^ mice. Similar treatments also normalized ERK1/2 phosphorylation **(Fig. 2L)** and surface AMPARs levels (Fig. S2D-E) in *Syngap1*^*PBM/PBM*^ neurons.

**Figure 2.**
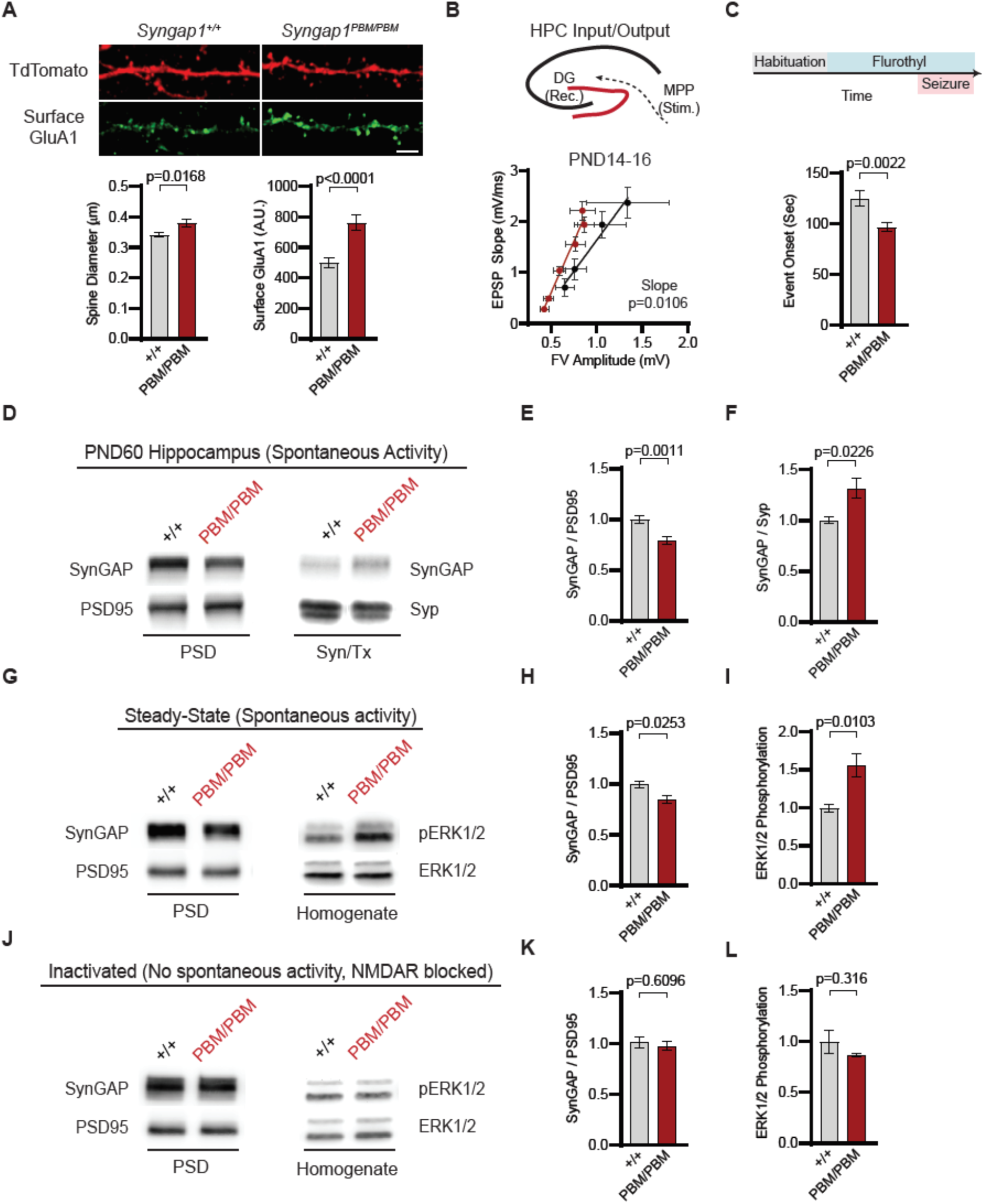
Disrupting the SynGAP PBM alters synapse properties downstream of NMDAR signaling. **(A)** Dendritic spine size and surface GluA1 expression levels in steady state primary forebrain cultures at DIV18. Average dendritic spine diameter. Unpaired t-test t(36)=2.508 p=0.0168. Average surface GluA1 expression in dendritic spines. t(36)=4.443 p<0.0001. n=19, each n representing an average of 25-30 spines fromge a dendritic segment belonging to distinct neuron. Scale bar 2 μm. **(B)** Hippocampal input-output relationships at PND 14-16 in dentate gyrus (DG). Linear regression comparing slopes and intercepts. Slope F(1, 6) = 13.36, p=0.0106, n=3 each n representing an individual mice (3-4 slices were recorded from each). **(C)** Seizure threshold measured as the time taken to reach the 1st clonus (event onset) during the procedure. Unpaired t-test. t(25)=3.420, p=0.0022, n=13-14 mice per genotype. **(D)** Western blots showing relative distribution of SynGAP in PSD and Syn/Tx fractions from adult hippocampi. **(E-F)** Quantification of western blots probing total SynGAP, Synaptophysin and PSD95. For PSD fractions PSD95 and for Syn/Tx fractions Synaptophysin (Syp) were used as loading control. PSD fractions: t(22)=3.733, p=0.0011 n=12 (3 technical replicates for each sample), Syn/TX fractions: t(6)=3.049, p=0.0226, n=4. Each sample represents hippocampi pooled from 2 mice. **(G)** Western blots showing relative enrichment of SynGAP and PSD95 in PSD fractions isolated from DIV18-21 cultures (left), phospho- and total-ERK1/2 levels in whole cell lysates in steady state (right). **(H-I)** Quantification of (G). Synaptic enrichment of SynGAP in steady state. Unpaired t-test: t(6)=2.961, p=0.0253, n=4. ERK1/2 phosphorylation is calculated as ratio of phospho-ERK1/2 to total-ERK1/2 in homogenates. t(12)=3.035, p=0.0103, n=6-8. **(J)** Western blots showing relative enrichment of SynGAP and PSD95 in PSD fractions (right), phospho and total-ERK1/2 levels in whole cell lysates in inactivated state (left). **(K-L)** Quantification of J. **(K)** Synaptic enrichment of SynGAP in inactivated state: t(6)=0.5385, p=0.6069, n=4. **(L)** ERK1/2 phosphorylation in inactivated state: t(4)=1.144, p=0.3163, n=3.

How does NMDAR activity interact with the PBM to regulate synaptic SynGAP expression and excitatory synapse signaling? First, because SynGAP PDZ binding has been shown to induce LLPS (Zeng et al., 2016), we explored the hypothesis that disrupting the PBM would alter the organization of macromolecular complexes within excitatory synapses. To test this, we immunoprecipitated PSD95 from neurons obtained from either WT or *Syngap1*^*PBM/PBM*^ mutant neurons. These neurons were treated with APV to avoid the confounds of elevated NMDAR signaling found in spontaneously active neurons from *Syngap1*^*PBM/PBM*^ mice **(Fig. 2A, D-F)**. These samples were then analyzed by mass spectrometry to determine how disrupting SynGAP-PDZ binding impacted the composition of PSD95 macromolecular complexes. In general, we found minor changes in the abundance of proteins that comprise PSD95 complexes in PBM mutants **(Fig. 3A)**. Only 1 (Gpm6a) out of ∼133 proteins known to be present within PSD95 complexes (Li et al., 2017) met our threshold for significance, although there were modest changes in proteins with structurally homologous PBMs (Type-1 PDZ ligands), such as Iqseq2 and Dlgap3 **(Fig. 3B)**. However, the vast majority of related PBM-containing proteins were not different in mutant neurons, including NMDAR subunits and TARPs (Fig. S3A). Consistent with the mass spectrometry analysis, immunoblot analyses found no changes in TARPs or LRRTM2 in isolated PSDs from *Syngap1*^*PBM/PBM*^ mice (Fig. S3B-E). Although PDZ binding was disrupted, SynGAP protein levels were also unchanged within PSD95 complexes, indicating that SynGAP may interact with PSD95 in a non-PDZ-dependent manner. There is significant overlap between the interactomes of PSD95 (Li et al., 2017) and SynGAP (Wilkinson et al., 2017) macromolecular complexes (Fig. S3F). Thus, within the intact postsynapse, SynGAP and PSD95 interact, as part of a macromolecular complex, through binding to common protein intermediaries. Together, these data suggest that SynGAP PBM binding to PDZ domains is not a major factor promoting the organization of PSD95 macromolecular complexes or the PSD.

**Figure 3.**
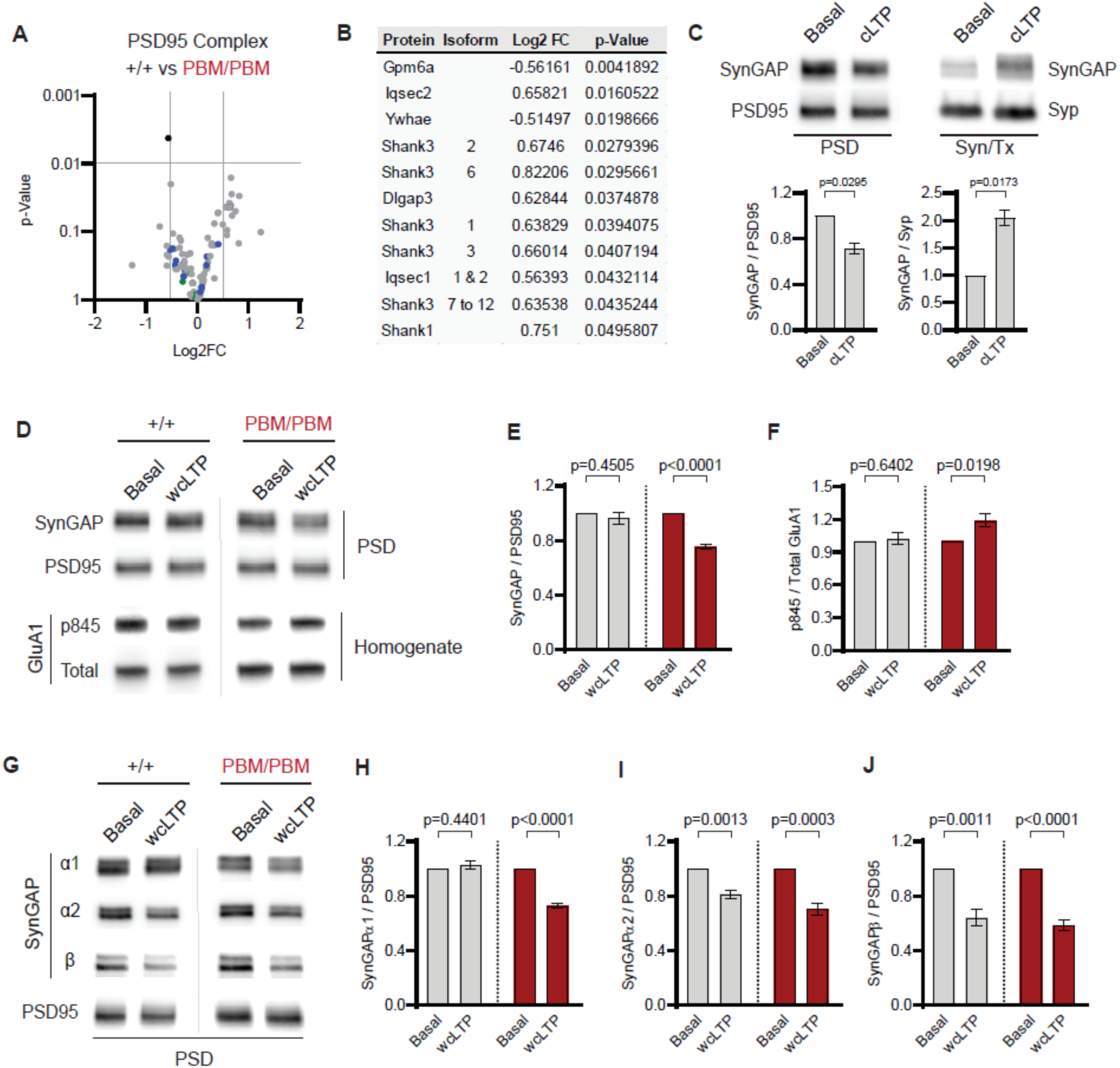
The SynGAP PBM increases the threshold for NMDAR signaling-dependent synaptic dispersion. **(A)** Volcano plot demonstrating the label-free quantitative mass-spectrometry profile of the logarithmic difference in protein levels in the immunoprecipitated PSD95 complexes derived from WT and PBM/PBM cultures in inactivated state. Only Gpm6a (shown in black) was significantly altered beyond the predetermined threshold (p<0.01 and ± 0.5 Log2 cutoff). Blue dots represent proteins with type 1 PDZ-ligands. Green dots represent DLG family proteins. P values were calculated via t-test for each protein. Samples were derived from individual cultures (4 per genotype) that were immunoprecipitated separately. **(B)** List of proteins differentially expressed beyond p>0.05 cutoff. Note that Iqseq2 and Dlgap3 are PDZ-binding proteins. **(C)** Blots showing distribution of SynGAP in PSD and Syn/Tx fractions upon chemical LTP stimulus (200μM Glycine, no Mg^+2^) in +/+ neurons. PSD: t(2)=5.696, p=0.0295. Syn/Tx: t(2)=7.511, p=0.0173. n=2, each n represents samples from separate cultures that are fractionated separately. **(D)** Representative blots from showing relative levels of SynGAP and PSD95 in PSD fractions or GluA1 Ser845 phosphorylation in basal state or weak chemical LTP (wcLTP: 10μM Glycine, no Mg^+2^) **(E-F)** Quantification of D. **(E)** Bar graphs represents the level of total-SynGAP/PSD95 ratio relative to the baseline of each genotype. +/+: t(4)=0.8354 p=0.4505, PBM: t(4)=17.52, p<0.0001, n=4 **(F)** GluA1 Ser845 phosphorylation. +/+: t(6)=0.4921 p=0.64, PBM: t(6)=3.152 p=0.0198 **(G)** Blots showing isoform enrichment in PSD fractions following wcLTP. **(H-J)** Quantification of G. **(H)** For SynGAP-α1 +/+: t(6)=0.8266 p=0.4401, PBM: t(6)=16.74 p<0.0001. **(I)** For SynGAP-α2, +/+: t(6)=5.706 p=0.0013, PBM: t(6)=7.263 p<0.0003. **(J)** For SynGAP-β, +/+: t(6)=5.894 p=0.0011, PBM: t(6)=10.52 p<0.0001, n=4. Note that α1 isoform is dispersed upon wcLTP only in PDZ-binding mutants.

We next explored the idea that the SynGAP PBM is important for SynGAP-specific dynamic functions, such as regulating it’s activity-dependent movement in and out of the PSD (Yang et al., 2011). As the synaptic phenotypes in PBM mutant mice were sensitive to APV treatments, we hypothesized that disrupting the SynGAP PBM would alter the sensitivity of activity-dependent signaling through NMDARs. To test this idea, we performed chemical LTP (cLTP) in WT and *Syngap1*^*PBM/PBM*^ mutant neurons. We confirmed that a standard cLTP procedure (Araki et al., 2015; Zeng et al., 2016), which has been shown to robustly activate synaptic NMDARs and promote excitatory synaptic plasticity, induced SynGAP extrusion in WT neurons **(Fig. 3C)**. However, a weak cLTP (Zeng et al., 2016), which results in reduced synaptic NMDAR activation, failed to induce t-SynGAP extrusion and GluA1 S845 phosphorylation in WT neurons **(Fig. 3D-F)**. In contrast, this weak stimulation was sufficient to trigger t-SynGAP extrusion from the PSD and induce GluA1 S845 phosphorylation in *Syngap1*^*PBM/PBM*^ neurons **(Fig. 3D-F)**, indicating that the threshold for signaling through NMDARs is regulated by the endogenous levels of SynGAP PBM at the PSD. As the PBM is splice-form specific, these data suggest that α1 binding to PSD-95 via its PDZ domains restricts its mobility out of the PSD in response to weak, sub-threshold NMDAR activation. To test this idea, we probed individual SynGAP isoforms in response to weak cLTP in WT and *Syngap1*^*PBM/PBM*^ neurons. Weak stimulation was sufficient to disperse the α1 isoform only in *Syngap1*^*PBM/PBM*^ neurons **(Fig. 3G,H)**, supporting the idea that PDZ binding limits NMDAR-dependent mobility of SynGAP. If PDZ binding limits isoform mobility during weak synaptic activation, then the SynGAP C-terminal isoforms that lack a PBM may be more sensitive to NMDAR activation (i.e. more likely to leave the PSD in response to a low level of NMDAR activity). To test this, we measured β and α2 isoform mobility in response to weak cLTP. In WT neurons, both isoforms were already dispersed from the PSD in response to weak cLTP **(Fig. 3G,I,J)**. Similar dynamics of these isoforms were observed in *Syngap1*^*PBM/PBM*^ neurons **(Fig. 3G,I,J)**, demonstrating that the PSD dynamics of α2 and β isoforms are regulated independently of α1. Given that t-SynGAP levels were unchanged in the PSD from WT neurons in response to weak cLTP **(Fig. 3D-E)**, our data indicate that t-SynGAP dynamics during synaptic NMDAR stimulation of primary neurons are dominated by α1. This may be a consequence of relative expression levels of SynGAP C-terminal isoforms within the PSD. Indeed, a recent study has suggested that α1 is the principle PSD-targeted SynGAP isoform in developing neurons (Gou et al., 2019).

NMDAR signaling is critical for hippocampal LTP and memory. Given that altering the SynGAP PBM disrupts signaling through NMDARs, we hypothesized that LTP would be disrupted in *Syngap1*^*PBM/PBM*^ mice. The within-train facilitation of responses across the seven theta bursts used to induce LTP did not differ between genotypes **(Fig. 4A)**, indicating that standard measures of induction, including NMDAR channel activation, were not impacted by PBM mutations. However, short-term plasticity (STP; **Fig. 4B, D)** and LTP **(Fig. 4C, E)** were both reduced in *Syngap1*^*PBM/PBM*^ mice. The ratio of LTP/STP was no different between genotypes **(Fig. 4F)**. Blocking NMDAR channel function disrupts both STP and LTP (Volianskis et al., 2013). However, a key measure of NMDA channel function was normal in PBM mutant mice **(Fig. 4A)**. Thus, these data are consistent with the idea that disrupting SynGAP-PDZ binding impairs signaling normally induced downstream of synaptic NMDAR activation.

**Figure 4.**
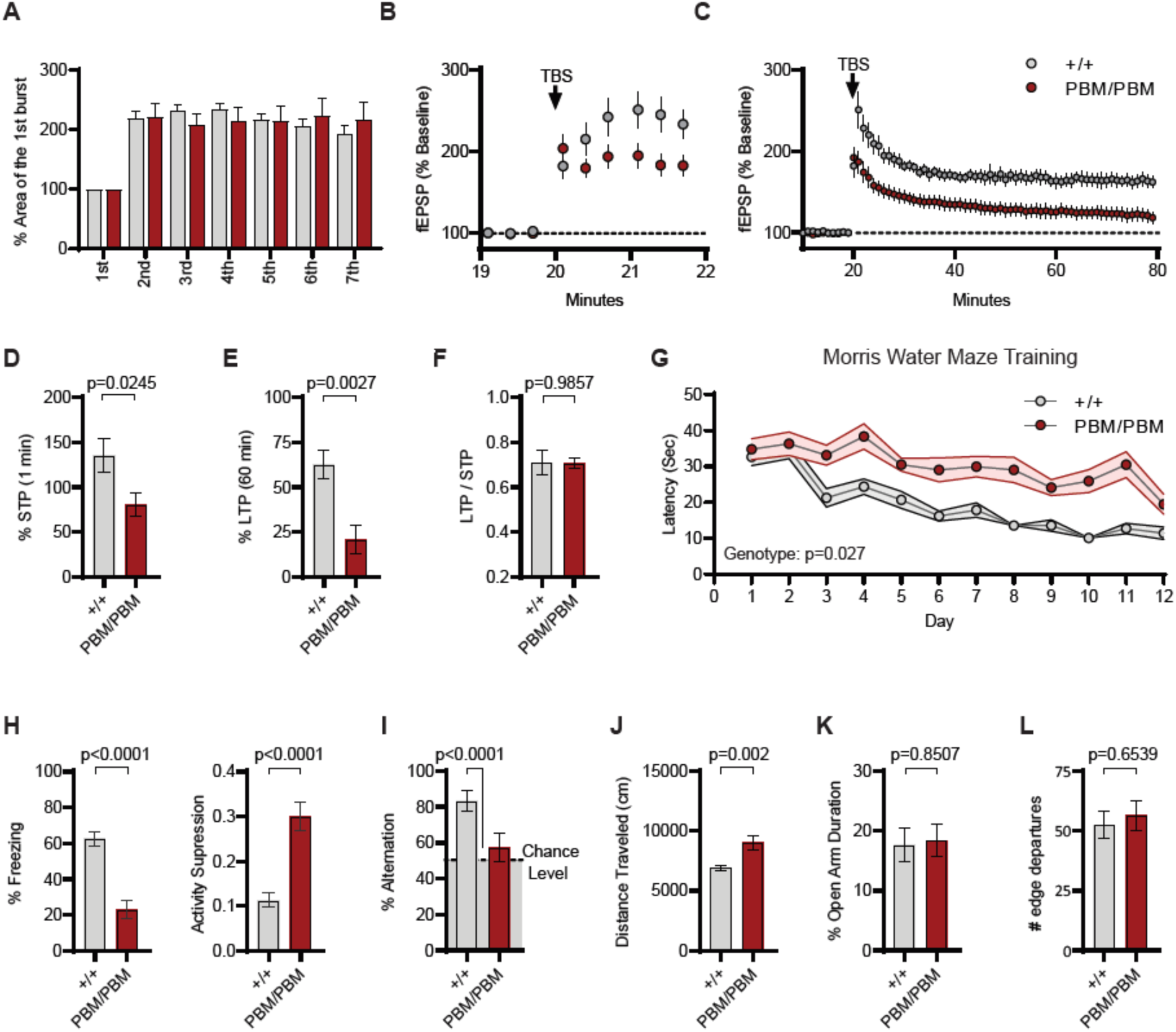
Loss of SynGAP-α1 PDZ-binding disrupts hippocampal synaptic plasticity and behaviors reflective of cognitive function: **(A)** Facilitation of burst responses calculated by expressing the area of the composite fEPSP corresponding to the 2nd theta burst within each train as a fraction of the 1st burst response. Genotype: F(1,84)=0.00034, p=0.9532. Burst number: F(5,84)=0.1448, p=0.9811, Interaction: F(5,84)=0.3065, p=0.9077. +/+: n=6, PBM/PBM: n=10. **(B)** Percent fEPSP during and immediately after the LTP induction. Note that homozygous mutants reach their peak potentiation immediately after TBS. **(C)** Magnitude of long-term potentiation (LTP) following delivery of a single train of seven theta bursts. The initial fEPSP slope was normalized to the mean value for a 20 min baseline period. **(D)** Graph shows short term potentiation (STP) % potentiation 1 min after TBS. t(15)=2.499, p=0.0245 **(E)** Graph shows % LTP 60 min after TBS. t(15)=3.594, p=0.0027 **(F)** LTP to STP ratio for individual slices. Note that the level of LTP is proportional to the degree of acute potentiation (1min post-TBS). t(15)=0.01818, p=0.9857. **(G)** Plots demonstrating latency to find platform across days in Morris Water Maze training session. Statistical significance was determined by using linear mixed model for repeated measures. Genotype: F(1,96)=5.066, p=0.027, n=14. **(H)** Percent freezing (left) and activity suppression (right) in remote contextual fear memory paradigm. % Freezing: t(45)=6.463, p<0.0001. Activity suppression: t(45)=5.599, p<0.0001 **(I)** Percent alternation in spontaneous alternation task measured via T-maze. One-sample t-test with 50% chance level as hypothetical mean were performed with each group for spontaneous alteration test. +/+ vs chance: t(25)=6.094, p<0.0001, n=26. PBM/PBM vs chance: t(20)=0.9005, p=0.3786, n=21. **(J)** Quantification of total distance traveled in the open field test (OFT). t(45)=3.427, p=0.0013. **(K)** Quantification of % time spent on open arm in elevated plus maze. t(35)=0.1896, p=0.8507. **(L)** Number of edge departures during the platform edge departure test. t(30)=0.4528, p=0.6539

Synaptic plasticity, such as LTP, is thought to contribute importantly to multiple forms of learning and memory. As such, we next measured performance of WT and *Syngap1*^*PBM/PBM*^ mice in a variety of learning and memory paradigms that have previously shown sensitivity in *Syngap1* knockout mouse models. We found evidence of impaired spatial learning in the Morris Water Maze **(Fig. 4G)** (Komiyama et al., 2002), disrupted associative memory in a remote contextual fear conditioning test **(Fig. 4H)** (Creson et al., 2019; Ozkan et al., 2014) and altered working memory during spontaneous alternation **(Fig. 4I)** (Clement et al., 2012; Muhia et al., 2010), in *Syngap1*^*PBM/PBM*^ mice. Moreover, we also observed increased horizontal locomotion in the open field test **(Fig. 4J)** (Guo et al., 2009; Muhia et al., 2010). There were no differences between genotypes in the elevated plus maze **(Fig. 4K)** and the cliff avoidance **(Fig. 4L)** tests, which are innate behaviors that are disrupted in *Syngap1* KO mice (Kilinc et al., 2018). Together, these behavioral data indicate that PBM-dependent mechanisms are critical for cognitive functions related to learning and memory, but less important for innate behavioral responses, such as risk-assessment and anxiety-related processes.

## Discussion

The main finding from this study is that the SynGAP PBM is critically important for regulating specific dynamic functions of SynGAP within the synapse, with implications for behaviors that reflect learning and memory. We genetically disrupted the endogenous SynGAP PBM to precisely understand how this motif contributes to synapse biology and behavior. Here, we demonstrate that a critical function of PBM binding is to limit the mobility of SynGAP-α1 in response to synaptic NMDAR activation. We found that weak NMDAR stimulation that does not reach a threshold for induction of synaptic potentiation is insufficient to eject the PBM-containing isoform, SynGAP-α1, from the PSD. This weak stimulation did, however, drive the removal of other, non-PBM-containing C-terminal isoforms (i.e. SynGAP-β and SynGAP-α2). Importantly, genetically disrupting the α1-specific PBM enhanced mobility of this isoform during weak stimulation, rendering its mobility similar to that of other non-PBM-containing isoforms. Removing PDZ binding of SynGAP-α1 had significant impacts on excitatory synapses. Neurons derived from PBM-deficient mice displayed less t-SynGAP at the PSD concomitant with elevated ERK1/2 signaling, enhanced AMPAR surface expression, and enlarged dendritic spines, as found in *Syngap1* conventional knockout mice (Kilinc et al., 2018; Rumbaugh et al., 2006; Vazquez et al., 2004). NMDAR blockade resulted in normal PSD levels of t-SynGAP and the rescue of these synaptic phenotypes. Together, the most parsimonious explanation for these results is that removing PDZ binding from α1 lowers the threshold for inducing NMDAR-dependent, Hebbian-type synaptic plasticity. In support of this idea, disrupting the endogenous PBM in SynGAP-α1 enabled weak chemical LTP to activate signaling pathways required for synaptic strengthening. Thus, PBM-mediated restriction of SynGAP-α1 mobility may help to set the threshold for NMDAR-dependent plasticity at excitatory synapses (Fig. S4). Our data indicate that a critical level of NMDAR activation is required to disperse α1 from the PSD. While inclusion of a PBM imparts SynGAP with unique features to regulate activity-dependent synaptic processes, the existence of a biochemical mechanism that dynamically regulates PBM affinity remains unknown. While this topic is beyond the scope of the current study, post-translational modifications of the SynGAP C-terminus, or the PDZ domain itself, may underlie such a mechanism. For example, phosphorylation of PBMs and/or a corresponding PDZ domain alters their affinity for each other (Lee and Zheng, 2010). SynGAP protein is a major target for kinases activated downstream of NMDARs, including CAMKII, CDK5, and PLK2 (Walkup et al., 2018; Walkup et al., 2015). If modifications of the SynGAP C-tail modulated PBM-PDZ affinity, then this could, in theory, enable dynamic plasticity thresholds at individual synapses, increasing the computational capacity within neural circuits. Indeed, we found in this study that modifying PBM binding affinity by genetically disrupting the α1 C-tail was sufficient to alter behaviors reflective of cognitive processes, such as learning and memory.

These results also have implications for disease. *De novo* pathogenic *SYNGAP1* variants in humans cause a relatively homogenous neurodevelopmental disorder defined by epileptic encephalopathy, intellectual disability, and autistic features (Deciphering Developmental Disorders, 2015, 2017; Hamdan et al., 2009; Mignot et al., 2016; O’Roak et al., 2014; Parker et al., 2015; Satterstrom et al., 2019; Vlaskamp et al., 2019). Our mouse model targeted a specific function of SynGAP-α1 (i.e. PDZ binding), with these genetic modifications causing significant impacts on synapse function and disease-relevant behaviors. Indeed, neurons from mice expressing a disrupted PBM displayed synapse phenotypes similar to what has been found in SynGAP haploinsufficient neurons (Kilinc et al., 2018), including enhanced synaptic excitability, larger dendritic spines, and impaired LTP. Moreover, behavioral phenotypes in PBM-deficient mice also phenocopied key aspects of the *Syngap1* heterozygous knockout mouse endophenotype, particularly with respect to behaviors reflective of impaired cognition and seizure (Kilinc et al., 2018). While there are no known *SYNGAP1* patient variants found in any of the final few exons of the gene that give rise to the major C-terminal isoforms (Jimenez-Gomez et al., 2019), our results imply that restoring function or expression of SynGAP-α1 could be therapeutically beneficial. Nearly all *SYNGAP1*-related NDD cases are thought to be caused by genetic haploinsufficiency (Weldon et al., 2018), or reduced SynGAP protein expression within brain cells. Therefore, *SYNGAP1* neurodevelopmental disorders are an attractive target for emerging therapies to restore gene function in patients. Gene restoration techniques, such as antisense oligonucleotide (ASO) injections or viral mediated gene replacement, have immense therapeutic potential (Sumner and Crawford, 2018). If these approaches are applied to *SYNGAP1* haploinsufficiency, our data suggest that upregulation of SynGAP-α1 would be beneficial. However, additional research is required to determine how re-expression of individual SynGAP C-terminal isoforms in animal models impact reversible phenotypes associated with the human disorder (Aceti et al., 2015; Clement et al., 2012; Creson et al., 2019; Ozkan et al., 2014).

## Acknowledgements

This work was supported in part by NIH grants from the National Institute of Mental Health (MH096847 and MH108408 to G.R., MH115005 and MH113949 to M.P.C, and MH105400 to C.A.M.), the National Institute for Neurological Disorders and Stroke (NS064079 to G.R.), the Eunice Kennedy Shriver National Institute of Child Health and Human Development (HD089491 to G.L.), the National Institute for Drug Abuse (DA034116 and DA036376 to C.A.M.), the Spanish Ministerio de Ciencia, Innovación y Universidades (BFU2012-34398, BFU2015-69717-P, RTI2018-097037-B-100, RYC-2011-08391p and IEDI-2017-00822) and the Catalan Government (AGAUR SGR14-297 and 2017SGR1776). M.K. was supported by Autism Speaks Weatherstone Pre-Doctoral fellowship (10646). G.G. was supported by a predoctoral fellowship from the Spanish Ministerio de Educación (BES-2013-063720).

## Author Contributions

M.K. performed experiments, designed experiments, analyzed data, co-wrote the manuscript, and edited the manuscript. T.K.C. performed experiments, designed experiments, analyzed data, and edited the manuscript. C.R. performed experiments, designed experiments and analyzed data. S.M. performed experiments, designed experiments, and analyzed data. A.A.L performed experiments, designed experiments, and analyzed data. J.L designed experiments and analyzed data. B.W. performed experiments, designed experiments, and analyzed data. N.H. performed experiments and designed experiments. N.G. performed experiments and designed experiments. A.R. analyzed data. G.G. performed experiments. Y.A. performed experiments. A.B. designed experiments, analyzed data, and interpreted data. M.P.K. designed experiments, analyzed data, and interpreted data. G.L. designed experiments, analyzed data, and interpreted data. C.A.M. designed experiments, interpreted data, and edited the manuscript. G.R. conceived the study, designed experiments, interpreted data, co-wrote the manuscript, and edited the manuscript.

## Materials & Methods

### Mice

*Syngap1* PBM mice were constructed in collaboration with genOway (France). The targeting vector (described in Fig. 1F) was electroporated into ES cells derived from the inner cell mass of 3.5 days old C57BL/6N embryos. Cells were then subjected to negative and/or positive selection(s) before the presence of the correct recombination event was validated by PCR and Southern blot. ES cell clones with verified mutations were injected into blastocysts which were implanted into pseudo-pregnant females to obtain chimeras. Chimeric mice were bred with C57BL/6 Cre-deleter mice to excise the Neomycin selection cassette and to generate heterozygous mice carrying the Neo-excised knock-in allele. Progeny was genotyped by PCR. The recombinase-mediated excision event was further validated by Southern blot using 5’ external probe. Knock-in lines were maintained on C57BL/6J background and bred for 3 generations prior to experimental use. Mice were genotyped using the following primers which amplified the locus spanning the LoxP site: Fwd: 5’-ctggttcaaaggctcctggta-3’ Rev: 5’-ctgtttgtttctcacctccaggaa-3’. This combination yielded a 61bp product in WT and 120bp product in knock-in alleles (Fig. 1H).

### Transcriptomics

PND7 mice forebrains (Cortex + hippocampus) were immediately removed and stored in RNALater (Thermo, AM7020). mRNA was isolated with RNeasy mini kit (74104, Qiagen). RNA integrity wass measured using Agilent 2100 Bioanalyzer (RIN value >= 9.2 for each sample). Library preparation and sequencing on the Illumina NextSeq 500 were performed by the Scripps Florida Genomics Core. De-multiplexed and quality filtered raw reads (fastq) were trimmed (adaptor sequences) using Flexbar 2.4 and aligned to the reference genome using TopHat version 2.0.9 (Trapnell et al., 2009). HT seqcount version 0.6.1 was used to generate gene counts and differential gene expression analysis was performed using Deseq2 (Anders and Huber, 2010). DeSeq2 identified differentially expressed genes (DEGs) with a cutoff of 1.5 fold change and an adjusted p-value of less than 0.05 (Love et al., 2014). Paired end reads mapped to the first 30 bases of Exon21 was used to determine the ratio of Exon21a (results in SynGAP-α2) vs Exon21b (results in SynGAP-α1) splicing events.

### Cell Culture

#### Cell lines

HeLa Cells (Kind gift of Michael Farzan) and HEK293T Cells (Kind gift of Joseph Kissil) were cultured in DMEM media containing 10% fetal bovine serum and penicillin/streptomycin.

#### Primary forebrain cultures

Dissociated forebrain cultures were prepared from newborn WT and homozygous littermates of the PBM line as previously described (Bedouin 2012). Briefly, forebrains were isolated and incubated with a digestion solution containing papain for 25 min at 37 °C. Tissues were washed and triturated in Neurobasal medium containing 5% FBS. Cells were plated on poly-D-lysine at a density of 1,000 cells per mm^2^. Cultures were maintained in Neurobasal A media (Invitrogen) supplemented with B-27 (Invitrogen) and Glutamax (Invitrogen). At DIV4 cells were treated with FuDR to prevent glial expansion. The cells were sparsely labeled by administration of AAVs (CamKII.Cre, 10^4^vg/ml, Addgene # 105558-AAV9 and CAG.Flex.EGFP, 10^8^vg/ml, Addgene #28304-PHPeB) at DIV 9-10 and processed for experiments 10-11 days later.

### *In situ* Colocalization Assay

HeLa cells were plated on glass coverslips and transfected with PSD95-tRFP (Plasmid #52671, Addgene) and/or EGFP-tagged SynGAP C-terminal constructs (EGFP-CCα1 or EGFP-CCPBM plasmids (made in house)) were co-transfected into HeLa cells using lipofectamine 2000 according to manufacturer’s instructions. Cells were then fixed with 4% PFA and washed multiple times with PBS prior to mounting with Prolong Gold with DAPI (P36931, Thermo). Confocal stacks spanning entire cells were obtained using UPlanSApo 100× 1.4 NA oil-immersion objective mounted on Olympus FV1000 laser-scanning confocal microscope using Nyquist criteria for digital imaging. Maximum intensity projections were used for the analysis. Nuclei of cells were defined by DAPI staining, and the EGFP-CC nuclear localization was calculated as the EGFP (colocalized with nucleus) / EGFP (within entire cell perimeter).

### PSD95-SynGAP Co-IP Assay

PSD95-tRFP (Plasmid #52671, Addgene) and/or full length EGFP-SynGAPα1/PBM (made in house) plasmids were transfected in HEK293T cells using Lipofectamine 2000. Cells were homogenized with Pierce IP Lysis buffer (87787, Thermo) containing protease & phosphatase inhibitors. Lysates were then incubated for 2hrs at RT with 1.5mg Dynabeads (10004D, Thermo) functionalized with 10ug of anti-PSD95 (Thermo, MA1-045) or IgG control (ab18415, Abcam). After extensive washing, immunoprecipitated proteins were eluted with Leammeli buffer at 70C for 10min with agitation. Eluted proteins were detected via western blot using PSD-95 (Thermo, MA1-045) and SynGAP (D20C7, CST) antibodies.10% of the input and 20% of IP elute were used for each sample.

### *In Vitro* Treatments

To silence neuronal activity and block NMDAR signaling, cultures were treated for 3hrs with 1 μM TTX and 200 μM APV. To induce chemical LTP, Cells were thoroughly washed and perfused with basal ECS (143 mM NaCl, 5 mM KCl, 10 mM HEPES (pH 7.42), 10 mM Glucose, 2 mM CaCl_2_, 1 mM MgCl_2_, 0.5 μM TTX, 1 μM Strychnine, and 20 μM Bicuculline) for 10 min. Then magnesium free ECS containing 200 μM Glycine (or 10 μM Glycine for weak cLTP) was applied for 10 min. Cells were then washed with and incubated in basal ECS for additional 10 min prior to downstream application.

### Subcellular Fractionation

#### From tissue

Frozen hippocampi or cortex were homogenized using a Teflon-glass homogenizer in ice-cold isotonic solution (320 mM sucrose, 50 mM Tris pH 7.4, phosphatase & protease inhibitors). The homogenate was then centrifuged at 1,000g for 10min at 4 °C. The supernatant (S1) was centrifuged at 21,000g for 30min. The pellet (P2) was resuspended in isotonic buffer and layered on top of a discontinuous sucrose density gradient (0.8M, 1.0M or 1.2M sucrose in 50mM Tris pH 7.4, +inhibitors) and centrifuged at 82,500g for 2hr at 4°C. The interface of 1.0M and 1.2M sucrose was collected as a synaptosomal fraction. Synaptosomes were diluted using 50mM Tris pH7.4 (+inhibitors) to bring the sucrose concentration to 320mM. The diluted synaptosomes were then pelleted by centrifugation at 21000g for 30min at 4°C. The synaptosome pellet was then resuspended in 50mM Tris pH 7.4 and then mixed with an equal part 2% Triton-X (+inhibitors). This mixture was incubated at 4 °C with rotation for 10min followed by centrifugation at 21,000xg for 20min to obtain a supernatant (Syn/Tx) and a pellet (PSD).

#### From primary culture

Cultured neurons (DIV 18-21), were homogenized by passage through 22G needle 10 times in ice-cold isotonic buffer (320 mM sucrose, 50 mM Tris, protease & phosphatase inhibitor mix). Homogenates were centrifuged at 1,000 × *g* for 10 min at 4 °C. The supernatant (S1) was centrifuged at 15,000 × *g* for 20 min at 4 °C to obtain the crude membrane (P2 fraction). The P2 pellet was resuspended with ice-cold hypotonic buffer (50 mM Tris, protease & phosphatase inhibitor mix) and was incubated for 30 min at 4C. Then the sample was centrifuged 21,000 x g for 30min to obtain synaptic plasma membrane (SPM) fraction. SPM is reconstituted in hypotonic buffer then equal volume of hypotonic buffer with 2% Triton-X was added and the mixture was incubated 15min on ice. Lysates were centrifuged at 21,000*g* for 30 min at 4 °C to obtain a soluble fraction (Syn/Tx) and a pellet (PSD), which was resuspended in 50 mM Tris containing 0.5% SDS. To completely solubilize PSD fraction, we’ve briefly sonicated and heated samples to 95 °C for 5min.

### Immunoblotting

Protein lysates were extracted from the hippocampi or cortices of adult mice and dissected in ice-cold PBS containing Phosphatase Inhibitor Cocktails 2 and 3 (Sigma-Aldrich, St. Louis, MO) and Mini-Complete Protease Inhibitor Cocktail (Roche Diagnostics) and immediately homogenized in RIPA buffer (Cell Signaling Technology, Danvers, MA), and stored at −80 °C. Sample protein concentrations were measured (Pierce BCA Protein Assay Kit, Thermo Scientific, Rockford, IL), and volumes were adjusted to normalize microgram per microliter protein content. For phospho-protein analysis, *in vitro* cultures were directly lysed with laemmeli sample buffer, sonicated and centrifuged to minimize DNA contamination. 10 μg of protein per sample were loaded and separated by SDS-PAGE on 4-15 % gradient stain-free tris-glycine gels (Mini Protean TGX, BioRad, Hercules, CA), transferred to low fluorescence PVDF membranes (45 μm) with the Trans-Blot Turbo System (BioRad). Membranes were blocked with 5% powdered milk (BSA for phospho-proteins) in TBST and probed overnight at 4 °C with the following primary antibodies: Pan-SynGAP (Thermo, PA1-046), SynGAP-α1 (Millipore, 06-900), SynGAP-α2 (abcam, ab77235), SynGAP-β (Kind gift of Rick Huganir), PSD-95 (Thermo, MA1-045), Synaptophysin (Novus, NB300-653), pERK (CST, 9106), ERK (CST, 4696), GluA1 (Millipore, MAB2263), phospho-serine845 GluA1 (Millipore, AB5847), TARP (Millipore, Ab9876), LRRTM2 (Thermo Pierce, PA521097).

### Immunocytochemistry

*For SynGAP – PSD95 colocalization*, neurons were fixed in 4% PFA, 4% sucrose for 5 min at RT and treated with MetOH for 15min at −20°C. The cells were then washed with PBS and permeabilized in PBS 0.2% TritonX-100 for 10 min. Samples were then blocked for 1 hr and probed for SynGAP (D20C7, CST) and PSD95 (MA1-045, Abcam) overnight. After PBS washes, samples were probed with appropriate secondary antibodies for 1 hr in the dark at room temperature. The coverslips were then washed, mounted (Prolong Glass) and cured. Confocal stacks were obtained. For analysis, maximum intensity Z projection was obtained from each confocal image. Individual synapses were traced as PSD95 positive puncta selected using an arbitrary threshold which was kept constant across all images. Mean SynGAP and PSD95 signals were measured from individual synapses. *For surface GluA1 staining*, neurons were immediately fixed in ice-cold pH 7.2 4% PFA, 4% sucrose for 20 min on ice. Then, samples were washed three times with ice-cold PBS and blocked for 1 hr min in PBS containing 10% NGS. Cells were then incubated overnight with a primary antibody targeting the extracellular N terminus of GluA1 (MAB2263, Millipore) and then washed with 10% goat serum twice to remove excess primary antibody. After PBS washes, Alexa dye–conjugated secondary antibodies were added for 1 hr in the dark at room temperature. The coverslips were then washed, mounted (Prolong Glass) and cured. Surface GluA1 levels were measured from manually traced individual dendritic spines from maximum intensity Z projection images using EGFP channel (cell fill). All confocal stacks were obtained for 6–12 individual fields from multiple coverslips per culture with UPlanSApo 100× 1.4 NA oil-immersion objective mounted on Olympus FV1000 laser-scanning confocal microscope using Nyquist criteria for digital imaging. 40-80 μm stretch of the secondary dendrites in neurons with pyramidal morphology were imaged.

### PSD95 Immunoprecipitation and Mass Spectrometry

Harvested neurons were lysed in DOC lysis buffer (50 mM Tris (pH 9), 30 mM NaF, 5 mM sodium orthovanadate, 20 mM β-glycerol phosphate, 20 µM ZnCl_2_, Roche cOmplete, and 1% sodium deoxycholate). The lysate was then centrifuged at 35,000 RPM for 30 minutes at 4°C and lysate containing 1 mg of protein was incubated with 2 µg Psd95 antibody (Neuromab, catalog # 75-048) at 4°C overnight with rotation. The following day, IPs were incubated with Dynabeads protein G (Thermo Fisher Scientific, catalog # 10004D) for 2 hours at 4 degrees Celsius. IPs were then washed three times with IP wash buffer (25 mM Tris (pH 7.4), 150 mM NaCl, 1 mM EDTA, and 1% Triton X-100). Dynabeads were re-suspended in 2X LDS sample buffer and incubated at 95 degrees Celsius for 15 minutes for elution. The eluate was incubated with DTT at a final concentration of 1 mM at 56°C for 1 hour followed by a 45-minute room temperature incubation with Iodoacetamide at a final concentration of 20 mM.

Samples were loaded onto 4 – 12% Bis-Tris gels and separated at 135V for 1.5 hours. Gels were stained with InstantBlue (Expedeon, catalog # 1SB1L) to visualize bands. The heavy and light chains of Immunoglobulin were manually removed. Gels were then destained using 25% ethanol overnight. Gel lanes were cut, individual gel slices were placed into 96 well plates for destaining, and peptide digestion was completed at 37 degrees Celsius overnight. Peptides were extracted with acetonitrile, dried down, and then desalted using stage tips. All LC-MS experiments were performed on a nanoscale UHPLC system (EASY-nLC1200, Thermo Scientific) connected to an Q Exactive Plus hybrid quadrupole-Orbitrap mass spectrometer equipped with a nanoelectrospray source (Thermo Scientific). Samples were resuspended in 10uL of Buffer A (0.1% FA) and 2uL were injected. Peptides were separated by a reversed-phase analytical column (PepMap RSLC C18, 2 μm, 100 Å, 75 μm X 25 cm) (Thermo Scientific). Flow rate was set to 300 nl/min at a gradient starting with 3% buffer B (0.1% FA, 80% acetonitrile) to 38% B in 110 minutes, then ramped to 75% B in 1 minute, then ramped to 85% B over 10 minutes and held at 85%B for 9 minutes. Peptides separated by the column were ionized at 2.0 kV in the positive ion mode. MS1 survey scans for DDA were acquired at resolution of 70k from 350 to 1,800 m/z, with maximum injection time of 100 ms and AGC target of 1e6. MS/MS fragmentation of the 10 most abundant ions were analyzed at a resolution of 17.5k, AGC target 5e4, maximum injection time 65 ms, and an NCE of 26. Dynamic exclusion was set to 30 s and ions with charge 1 and >6 were excluded. The maximum pressure was set to 1,180 bar and column temperature was constant at 50°C.

Proteome Discoverer 2.2 (Thermo Fisher Scientific) was used to process MS data and analyzed using Sequest HT against Uniprot mouse databases combined with its decoy database. With respect to analysis settings, the mass tolerance was set 10 parts per million for precursor ions and 0.02 daltons for fragment ions, no more than two missed cleavage sites were allowed, static modification was set as cysteine carbamidomethylation, and oxidation of methionine was set as a dynamic modification. False discovery rates (FDRs) were automatically calculated by the Percolator node of Proteome Discoverer with a peptide and protein FDR cutoff of 0.01. Label free quantification was performed using Minora node in Proteome Discoverer. Abundances of identified PSD95 interacting proteins in WT and mutant neurons were compared using relative abundances such that proteins with a fold change in abundance ratio of > 2.0 or < 0.5 were considered to be differentially associated to PSD95.

### Hippocampal LTP and Extracellular Recordings

Acute transverse hippocampal slices (350 µm) were prepared using a Leica Vibroslicer (VT 1000S), as described previously (Babayan et al., 2012). Slices were cut into ice cold, choline chloride artificial cerebral spinal fluid (ACSF) containing (in mM) 110 choline chloride, 2.5 KCl, 1.25 NaH2PO4, 5 MgSO4, 25 NaHCO2, 25 glucose, 11.6 ascorbic acid, and 3.1 pyruvic acid and rinsed at room temperature for ∼3 min in a high magnesium aCSF solution containing: 124 NaCl, 3 KCl, 1.25 KH2PO4, 5 MgSO4, 26 NaHCO3, and 10 dextrose. Slices were then transferred to an interface recording chamber maintained at 31±1°C, oxygenated in 95% O2/ 5% CO2 and constantly perfused (60-80 ml/h) with normal ACSF (in mM; 124 NaCl, 3 KCl, 1.25 KH2PO4, 1.5 MgSO4, 2.5 CaCl2, 26 NaHCO3, and 10 dextrose). Slices equilibrated in the chamber for approximately 2 hours before experimental use. Field excitatory postsynaptic potentials (fEPSPs) were recorded from CA1b stratum radiatum using a single glass pipette (2-3 MΩ). Bipolar stainless-steel stimulation electrodes (25 µm diameter, FHC) were positioned at two sites (CA1a and CA1c) in the apical Schaffer collateral-commissural projections to provide activation of separate converging pathways of CA1b pyramidal cells. Pulses were administered in an alternating fashion to the two electrodes at 0.05 Hz using a current that elicited a 50% maximal response. After establishing a 10-20 min stable baseline, long-term potentiation (LTP) was induced in the experimental pathway by delivering 7 ‘theta’ bursts, with each burst consisting of four pulses at 100 Hz and the bursts themselves separated by 200 msec (i.e., theta burst stimulation or TBS). The stimulation intensity was not increased during TBS. The control pathway received baseline stimulation (0.05Hz) to monitor the health of the slice. The fEPSP slope was measured at 10–90% fall of the slope and all values pre- and post-TBS normalized to mean values for the last 10 min of baseline recording. Baseline measures for all groups included paired-pulse facilitation and input/output curves.

### Behavior

At weaning, four mice were randomly allocated to one cage with respect to genotype with males and females being housed separately. Randomization of cage allocation was restricted in that, as much as possible, mice from the same litter were placed in different cages so that no single litter was overrepresented in any single experiment. Cages utilized for behaviors contained cardboard pyramidal-shaped huts with two square openings on opposing sides of the hut for the purposes of environmental enrichment and to assist with transfers from home cages to behavioral apparatuses. All mice were handled for several minutes on three consecutive days prior to commencement of behavioral testing. Tails were marked for easy identification and access from home cages during testing. Experimenters were blind to mouse genotype while conducting all tests. All mice were backcrossed onto a C57BL/6J genetic background for at least 3 generations.

#### Flurothyl-induced seizures

Flurothyl-induced seizure studies were performed based on prior studies with some modifications (Clement et al., 2012; Dravid et al., 2007; Ozkan et al., 2014). Briefly, experiments were conducted in a chemical fume hood. Mice were brought to the experimental area at least 1 h before testing. To elicit seizures, individual mice were placed in a closed 2.4-L Plexiglas chamber and exposed to 99% Bis (2,2,2-triflurothyl) ether (Catalog# 287571, Sigma-Aldrich, St. Louis, MO). The flurothyl compound was infused onto a filter paper pad, suspended at the top of the Plexiglas chamber through a 16G hypodermic needle and tube connected to a 1 ml BD glass syringe fixed to an infusion pump (KD Scientific, Holliston, MA, USA, Model: 780101) at a rate of 0.25 ml/min. The infusion was terminated after the onset of a hind limb extension that usually resulted in death. Cervical dislocation was performed subsequently to ensure death of the animal. Seizure threshold was measured as latency (s) from the beginning of the flurothyl infusion to the beginning of the first myoclonic jerk.

#### Morris water maze

Mice were run in a standard comprehensive Morris water maze paradigm including a cue test with a visual platform and an acquisition protocol with a hidden platform. All phases of the paradigm were run in a dedicated water maze room in the Scripps Florida Mouse Behavior Core. A water maze system including a plastic white opaque pool (Cat# ENV-594M-W, Med Associates), measuring ∼122cm diameter at the water surface, supported by a stand (ENV-593M-C) and equipped with a floor insert (ENV-595M-FL) covering a submerged heater was utilized for all water maze experimentation. An adjustable textured platform (17.8 cm diameter, ENV-596M) was placed atop the floor insert in one of two different quadrants, depending on the specific phase of the paradigm (NW quadrant for initial training and probe test and SE quadrant for reversal training and probe tests), for mice to escape the water. Water temperatures were controlled to 22.5 ± 0.5 °C using a built-in heater and monitored with a digital temperature probe. This water temperature motivated the mice to escape the water without eliciting hypothermic conditions. The tank was emptied, cleaned and refilled once every three days to avoid unsafe accumulation of bacteria. Water was made opaque by the addition of a white opaque non-toxic paint (Crayola) forcing mice to utilize extra-maze cues when locating the hidden platform (0.5 cm beneath the surface of the water). These spatial cues (large black cardboard circle, star, square, white X on black background) were placed on the walls of the room at different distances from the pool. The pool edge was demarcated with directional units (W, N, E, S) to aid assignment of invisible platform “quadrants” to the pool arena outlined by the video tracking system. Various strip lights were positioned on the walls near the ceiling to allow for a moderate level of lighting (200 lux), enough for the mice to see the extra-maze cues adequately without eliciting undue anxiety. Thirty minutes prior to commencement of daily trials, the lights and heater were turned on, and mouse home cages were placed on heating pads on a rack in the water maze room to provide a warm place for the mice between trials. Cage nestlets were replaced with strips of paper towels to better facilitate drying after trials. Mice were monitored during trials for signs of distress and swimming competence. None of the mice tested had swimming issues, and floating was discouraged with gentle nudges. Mice received four trials per day during cue and acquisition phases and one trial per day for probe trials. Three cages (12 mice) were run at a time such that ITIs for each day lasted about 20 minutes with trial duration lasting until the mouse found the platform or a maximum of 60 s. Each trial commenced when the mouse was automatically detected in the pool by the tracking system (Ethovision, Noldus). Each mouse was lowered into the pool facing its edge at one of the four directional units (W, N, E, S) in a clockwise manner, with the first of the four trials starting closest to the platform (“NW quadrant”), which was positioned in the central area of the quadrant dictated by the tracking system. This same series of daily trial commencements were followed for all mice for each of the cue tests, acquisition protocol, and reversal protocol. If the mouse did not locate the platform in 60 s, the experimenter’s hand guided them to the platform. Because the mice are eager to escape the water, the mice quickly learned to follow hand direction to the platform, minimizing physical manipulation of the animals during the trials. Mice were allowed 15 seconds on the platform at the end of each trial before being picked up, dried with absorbent wipes, and placed back into their warmed home cage.

On the first day of testing, mice were given a cue test with the platform positioned just above the surface of the water and a metal blue flag placed upon it for easy visual location of the platform. This test allows for detection of individual visual and swimming-related motor deficits and allows the mice to habituate to the task (climbing on the platform to escape the water). The platform was placed in a different location for each of the four trials with spatial cues removed by encirclement of the pool with a white plastic curtain.

On the next day, acquisition trials began with the hidden platform remaining in the same location (“NW quadrant”) for all trials/days and the curtain drawn back for visibility of the spatial cues. Several measures (distances to platform) and criteria to reach the platform (approximately 90% success rate, approximately 20 second latency to find platform) during the acquisition phases were recorded and achieved before mice were deemed to have learned the task. The performances of the four trials were averaged for each animal per day until criteria were met.

#### Elevated plus maze

A dedicated OFT/Elevated Plus Maze (EPM) room in the Mouse Behavioral Core containing a black acrylic EPM apparatus (Model ENV-560A, Med Associates, St. Albans, VT) consisting of two closed arms and two open arms extending from an open square center area was used to measure anxiety-like or risk-taking behaviors. The arms of the maze (34.9 × 6 cm) were situated in a plus orientation with each of the open arms and each of the closed arms radiating 180 degrees from each other. Closed arm walls were 19.7 cm high extending from the base of the arm; open arms had no walls other than a 0.5 cm vertical edge. Thick white corrugated cardboard was fitted into the arm floors for contrast with the dark color of the mice for video recording purposes. The entire apparatus was placed in a far corner of the room with a two-walled construct at the opposite corner. A white noise generator (2325-0144, San Diego Instruments) was set at 65 dB to mask external noises and provide a constant noise level. Fluorescent linear strip lights placed on each of the four walls of the behavioral room adjacent to the ceiling provided a lower lighting (200 lux) environment than ceiling lighting to encourage exploration with closed arms receiving half the lighting levels as that of the open arms. One CCTV camera (Panasonic WV-BP334) was connected to a computer equipped with tracking software (Ethovision XT 11.5 (Noldus Information Technology, Leesburg, VA)). At the beginning of each trial, mice were individually placed in the center of the maze facing the open arm toward the far wall of the room. Each mouse was allowed to explore the maze for 5 min and analyzed for percent time spent in the open arms.

#### Open field test

Naive mice were individually introduced into one of eight adjacent open field arenas for 30 min and allowed to explore. Open field arenas consisted of custom made clear acrylic boxes (43 × 43 × 32h cm) with opaque white acrylic siding surrounding each box 45 × 45 × 21.5h cm to prevent distractions from activities in adjacent boxes. Activity was monitored with two CCTV cameras (Panasonic WV-BP334) feeding into a computer equipped with Ethovision XT 11.5 for data acquisition and analyses. A white noise generator (2325-0144, San Diego Instruments) was set at 65 dB to mask external noises and provide a constant noise level. Fluorescent linear strip lights placed on each of the four walls of the behavioral room adjacent to the ceiling provided a lower lighting (200 lux) environment than ceiling lighting to encourage exploration.

#### Platform edge departure test

The risk-taking/impulsivity test in adult *Syngap1* mice was carried out using young (∼PND21) mice. C57BL/6 wild type and mutant mice were weaned the day before the test was conducted. Each mouse, two at a time, was placed on an overturned stainless-steel wire Galaxy pencil/utility cup (7.8 cm bottom diameter (“platform”), 10.8 cm high) situated on a black granite bench top in a vivarium procedure room within demarcated areas (30×30×27 cm) using box cardboard dividers to separate the two mice from each other and two other sides on the bench with a front open side where a notebook web camera videotaped each 10 min session. White duct tape was affixed to the black foam bottom of both cups for easier cleaning between trials and better contrasts with black mice. Each mouse’s activities were hand scored thereafter. The number of “edge departures” was the principle measure. This behavior was defined by the mouse forepaws grasping the platform edge or wire mesh that made up the vertical walls exhibiting a forward stance over the edge. The latency to first full departure was also scored. Platform departures occurred when all four paws were the top platform and on the vertical bars. Pilot tests using other cups or beakers of different heights did not alter scoring parameter measures. However, we preferred the wire sided cups because we could measure intended edge departures as well as climbing tendencies not possible with a glass beaker eliminating most unintended loss-of-balance movements from the platform. Experimenters and scorers were blind to group identities with several individual scorers attaining similar parameter measurements.

#### Spontaneous alternation

A dedicated T-maze room in the Mouse Behavior Core containing four T-mazes placed in various orientations was used to assess working memory in a discreet-two-trials spontaneous alternation (SA) paradigm. Each maze contained three arms with walls made opaque including a start box (17.8 × 7.3 cm) at the base of the start arm (38.1 × 7.3 cm) and adjoined to a central choice area (10.2 × 10.2 cm) with two choice arms 30.5 × 7.3 cm) radiating 180 degrees from the central choice area. Automatic guillotine doors were installed at the end of the start arm box and at the entrances of each of the choice arms that were activated by motion detected by an Ethovision XT 11.5 system linked to CCTV cameras. Two of the mazes oriented 180 degrees from each other were used for all assessments with each mouse tested in a different maze on four consecutive days. Each test consisted of two free-choice trials separated by a 1 min inter-trial interval (ITI) when the mouse was confined to the start box. Mice were given one set of trials per day for four days. At the commencement of each test session, the mouse was placed in the start box with a guillotine door opening once the animal was detected. Once the mouse traveled down the start arm and into one of the choice arms that door closed. The mouse was left in this choice arm for 10 s before being gently scooped up and returned to the start arm for Trial 2. After the 1 min ITI, the start box door automatically opened, and the mouse was allowed another free-choice trial. If the mouse entered the opposite arm from that of the first trial, the responses were recorded as an alternation. If mice did not move from the start box within 3 min on Trial 1 or within 1 min on Trial 2, they were gently nudged. Mice were excluded from data analyses for the sessions they were uncooperative.

#### Contextual fear conditioning

A dedicated fear conditioning room in the TSRI Florida Mouse Behavior Core contains four fear conditioning devices that can be used in parallel. Each apparatus was an acrylic chamber measuring approximately 30 × 30 cm (modified Phenotyper chambers, Noldus, Leesburg, VA). The top of the chamber is covered with a unit that includes a camera and infrared lighting arrays (Noldus, Ethovision XT 11.5, Leesburg, VA) for monitoring of the mice. The bottom of the chamber is a grid floor that receives an electric shock from a shock scrambler that is calibrated to 0.40 mA prior to experiments. The front of the chamber has a sliding door that allows for easy access to the mouse. The chamber is enclosed in a sound-attenuating cubicle (Med Associates) equipped with a small fan for ventilation. Black circular, rectangular and white/black diagonal patterned cues were placed outside each chamber on the inside walls of the cubicles for contextual enhancement. A strip light attached to the ceilings of the cubicles provided illumination. A white noise generator (∼65 dB) was turned on and faced toward the corner of the room between the cubicles. The fear conditioning paradigm consisted of two phases, training, followed by testing 1 and 26, or 30 d thereafter. The 4.5 min training phase consisted of 2.5 min of uninterrupted exploration. Two shocks (0.40 mA, 2 s) were delivered, one at 2 min 28 s, the other at 3 min and 28 s from the beginning of the trial. During testing, mice were placed into their designated chambers and allowed to roam freely for 5 min. Immobility durations (s) and activity (distances moved (cm)) during training and testing were obtained automatically from videos generated by Ethovision software. Activity suppression ratio levels were calculated: 0-2 min activity during testing/0-2 min activity during training + testing.

