## Supplementary Figures for "Splicing of the SynGAP Carboxyl-Terminus Enables Isoform-Specific Tuning of NMDA Receptor Signaling Linked to Cognitive Function"

Figure S1

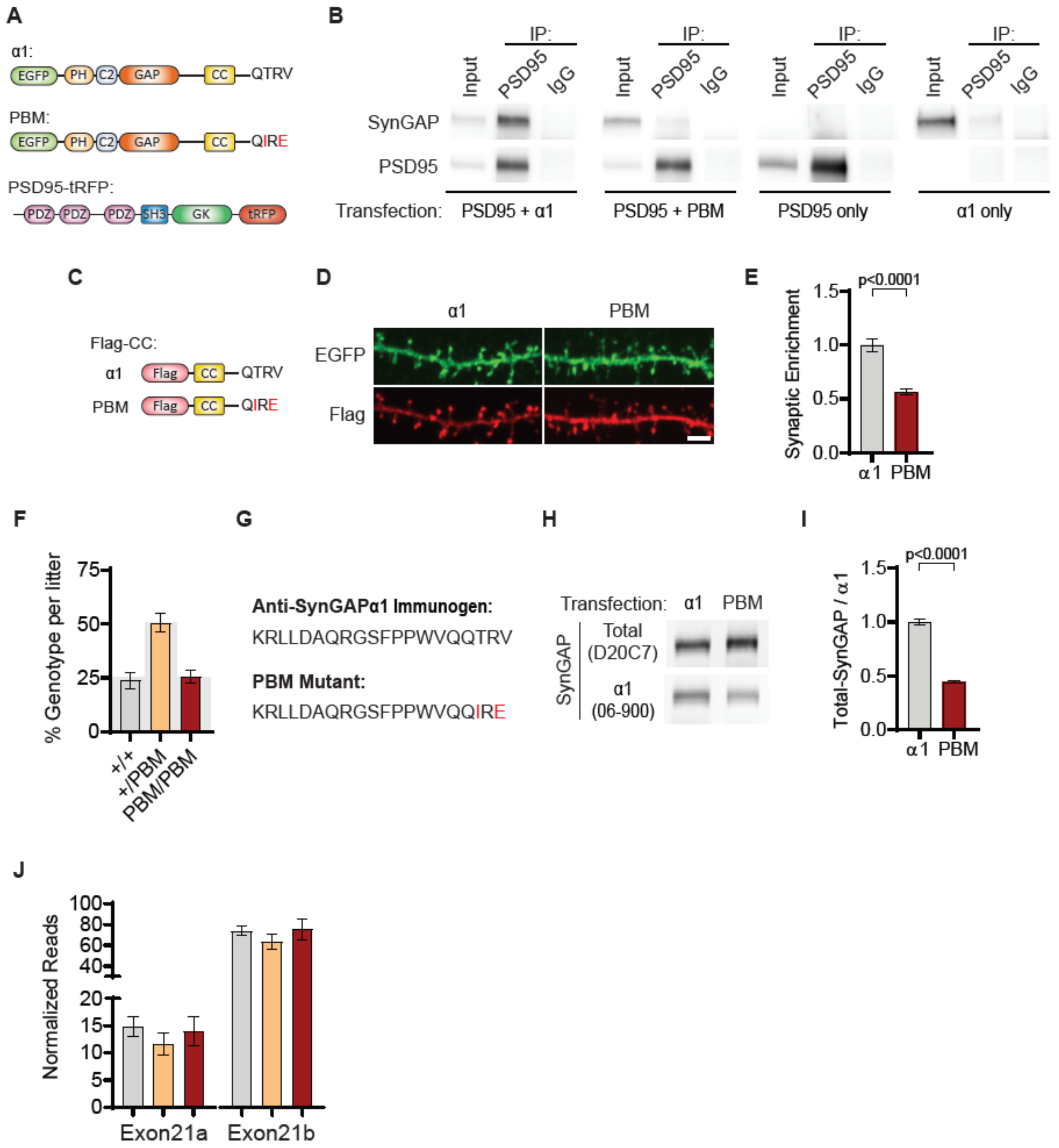


**Figure S1. (A)** Illustrations of constructs expressed in H293T cells to study PDZ-dependent interaction between SynGAP and PSD95. **(B)** Coimmunoprecipitation of PSD-95 and SynGAPα1 from transfected H293T cells. PSD95-tRFP coprecipitates with SynGAP-α1. This interaction was disrupted by PBM mutations. **(C)** Illustrations of Flag-tagged SynGAP C-terminal constructs expressed in primary cortical neurons. **(D)** Subcellular localization of wild-type or PBM mutated Flag-CCα1 in primary forebrain neurons. Note that Flag-CC-α1 is heavily enriched in dendritic spines compared to Flag-CCPBM. Scale bar 2µm. **(E)** Quantification of synaptic enrichment of Flag-CC constructs. Enrichment in dendritic spines were calculated as the ratio of Flag signal in spines vs dendrites over ratio of EGFP signal in spines vs dendrites. Unpaired t-test, $t(9)=6.982$ $p<0.0001$. **(F)** Genotype distribution of 15 litters (128 mice) produced by *Syngap1*$^{+/PBM}$ interbreeding. Expected genotype frequency is highlighted in gray. **(G)** Antigen sequence for α1-specific antibody in comparison to PBM mutant C-tail. **(H)** Western blots showing reduced antigenicity of α1 antibody against PBM mutant C-terminus. H293T cells were transfected with either wild-type or PDZ-binding mutant form of EGFP-SynGAP-α1. Lysates were probed for both Pan-SynGAP (D20C7) and α1-specific (06-800) antibody. Relative reduction in α1 to Pan-SynGAP signal demonstrates ~60% reduction in antigenicity. **(I)** Quantification of (H) Unpaired t-test. $t(6)=19.16$, $n=4$, $p<0.0001$. **(J)** SynGAP exon 21 splicing in forebrain transcriptome. Bar graph demonstrates normalized sequencing depth of paired end reads covering exon 21a (specific to α2) and exon 21b (specific to α1). For exon 21a, Genotype: $F(2,6)=0.5924$, $p=0.5824$. For exon 21b, Genotype: $F(2,6)=0.8049$, $p=0.4901$, $n=3$.

### Figure S2

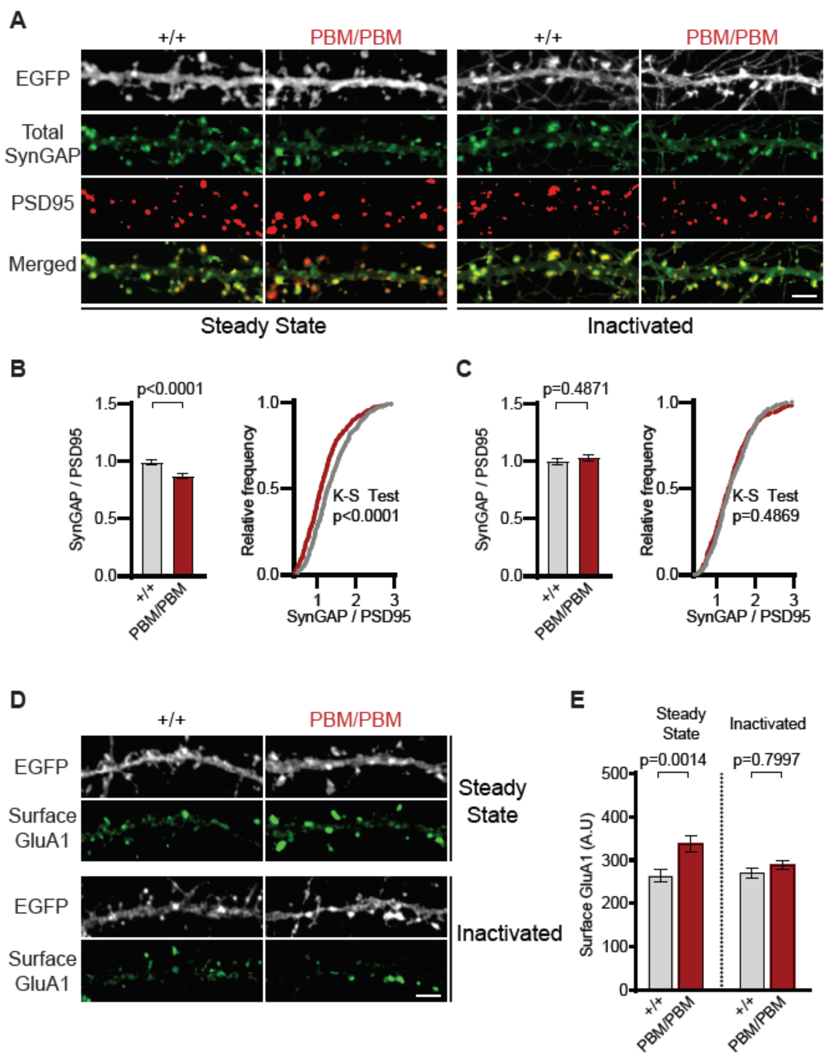


**Figure S2. (A)** Representative images showing synaptic enrichment of total SynGAP in WT and PBM mutants in primary neuron cultures in steady or inactivated state. Scale bar 2 μm. **(B, C)** Levels of SynGAP relative to PSD95 signal in dendritic spines. Left, bar graphs demonstrate mean enrichment in an individual dendritic segment. Steady state: t(90)=4.393, p<0.0001, n=45-47. Inactivated: t(78)=0.6982, p=0.48, n=38-44. Right, cumulative distribution of SynGAP to PSD95 ratios in individual synapses. Kolmogorov-Smirnov test, Steady state: p<0.0001, Inactivated: p=0.4869. **(D)** Representative images showing surface GluA1 expression in primary forebrain cultures in steady or inactivated state. Scale bar 2 μm. **(E)** Quantification of mean surface GluA1 levels coincident with PSD95 puncta. Two-way ANOVA with Tukey's multiple comparisons test. Interaction: F(1,74)=4.112, p=0.0462, Genotype: F(1,74)=11.09, p=0.0014.

Treatment: $F(1,74)=2.329$, $p=0.1313$. $n=19$-$21$, each n representing an average of 25-30 spines from a dendritic segment belonging to distinct neurons.

**Figure S3**

**A**

| Type-1 PDZ Ligand Containing Proteins | | |
|---|---|---|
| Gene | Log2FC | p-Value |
| Iqsec2 | 0.65821 | 0.016052246 |
| Dlgap3 | 0.62844 | 0.037487805 |
| Dlgap4 | 0.40918 | 0.051794849 |
| Dlgap1 | 0.33653 | 0.075018022 |
| Cyfip2 | 1.2431 | 0.100533928 |
| Dlgap2 | 0.28254 | 0.147590456 |
| Grin2d | 0.41014 | 0.151788442 |
| Dlgap2 | 0.27765 | 0.155441427 |
| Syngap1 | -0.48327 | 0.175547053 |
| Gria1 | -0.52362 | 0.181726733 |
| Adam22 | -0.58291 | 0.214676179 |
| Grin2a | 0.18881 | 0.229584435 |
| Cacng2 | -0.4177 | 0.262876952 |
| Ncam1 | -0.24765 | 0.279263266 |
| Grin2b | 0.18197 | 0.306477882 |
| Gda | -1.26898 | 0.307778237 |
| Btbd11 | -0.23161 | 0.333135481 |
| Dlgap1 | 0.19309 | 0.34394071 |
| Abr | 0.06289 | 0.457748635 |
| Dstn | -0.13564 | 0.822744192 |

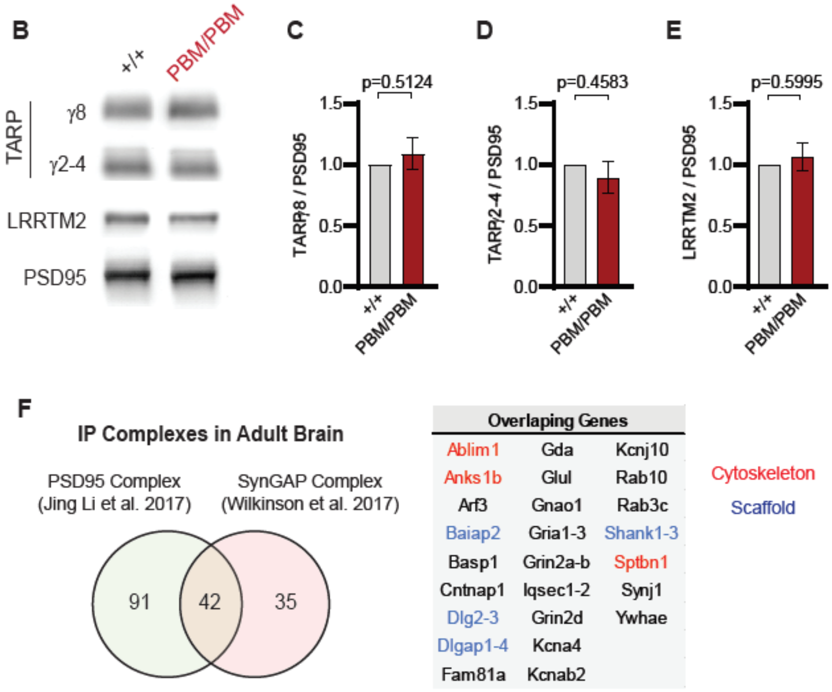


**Figure S3. (A)** Mass-spectrometry profile of type-1 PDZ binding motif containing proteins in immunoprecipitated PSD95 complex in +/+ vs PBM/PBM inactivated cultures. **(B)** Western blots showing relative expression of TARPs and Lrrtm2 in PSD fractions from adult hippocampi in +/+ vs PBM/PBM. **(C-E)** Quantification of B. **(C)** TARPγ8 t(6)=0.6961, p=0.5124. **(D)** TARPγ2-4 t(6)=0.7924, p=0.4583. **(E)** LRRTM2 t(6)=0.5542, p=0.5995, n=4, each representing hippocampi pooled from 2 mice. **(F)** Comparison of PSD95 and SynGAP IP complexes as reported by (Li et al. 2017) and (Wilkinson et al. 2017). Note that PSD95 and SynGAP complexes share diverse range of components involving cytoskeletal and scaffolding proteins.

### Figure S4

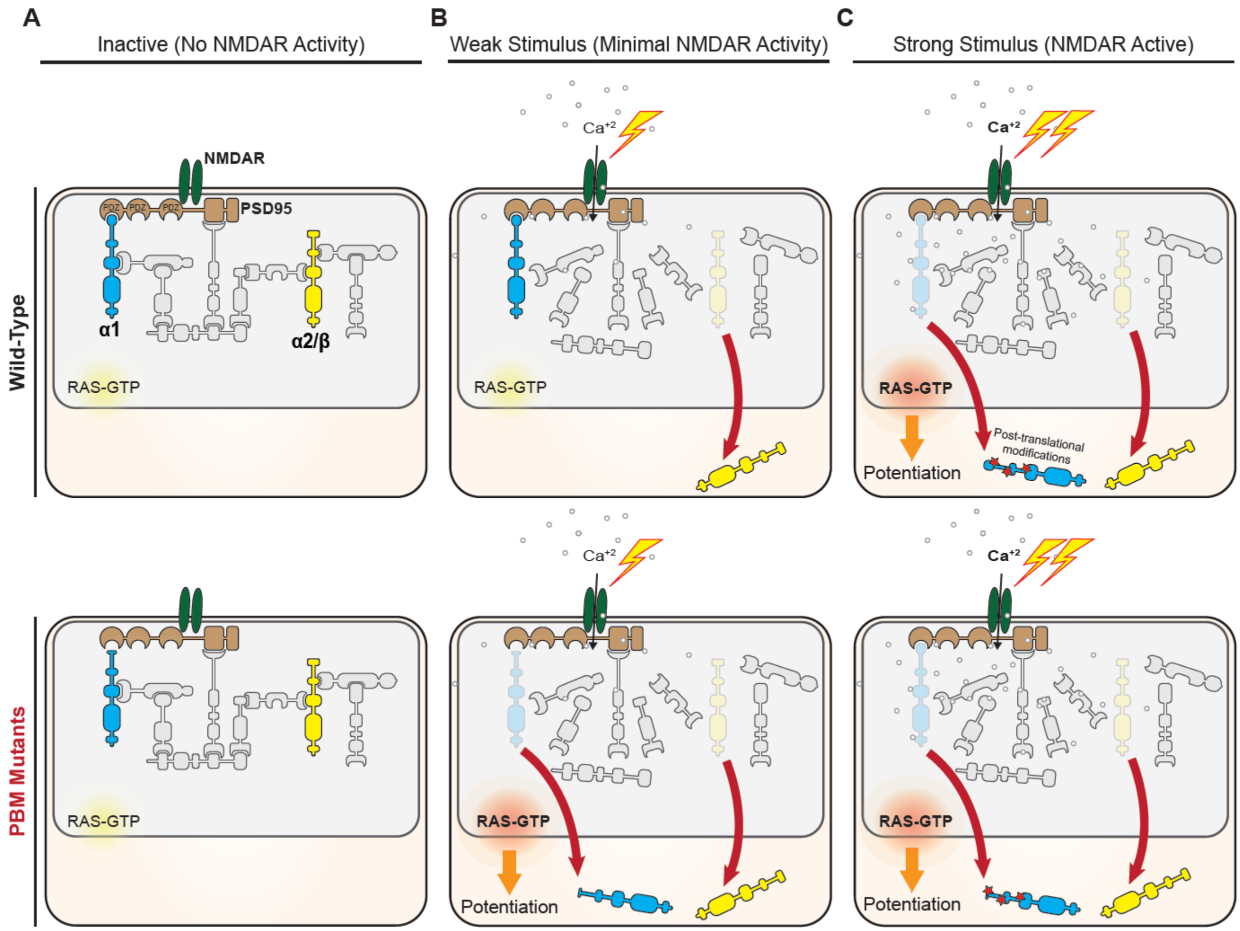


**Figure S4. (A)** In the absence of NMDAR activity, SynGAP serves as part of the PSD95 complex through PDZ-independent interactions. Isoforms devoid of PBM (α2/β or α1 PBM mutants) are still present in postsynaptic density. **(B)** In the presence of weak NMDAR activity, isoforms without PDZ binding are more likely to disperse out of the PSD. The PBM in α1 increases the affinity of this isoform to PSD, which reduces its PSD mobility following weak NMDAR activity. In PBM mutants, impaired PDZ-binding reverts α1 mobility similar to non-PBM isoforms. **(C)** Upon strong NMDAR activity, post-translational modifications (e.g., phosphorylation) of SynGAP may reduce PBM affinity for PDZ domains, which results in dispersion of α1 isoform. Because SynGAP is a RasGAP, the dynamic reduction of endogenous α1 from the PSD in response to strong NMDAR activation promotes GTPase signaling at dendritic spines. Thus, PBM binding within α1 serves as a molecular trigger, that when engaged, helps to converts NMDAR channel activity into dendritic spine GTPase signaling required for synaptic potentiation.
